# Hematopoietic growth factors Regulate Entry of Monocytes into the Adult Brain via Chemokine Receptor CCR5

**DOI:** 10.1101/2024.05.15.594359

**Authors:** Xuefang Ren, Junchi He, Heng Hu, Shinichi Kohsaka, Li-Ru Zhao

**Affiliations:** Department of Neurosurgery, State University of New York Upstate Medical University, Syracuse, New York 13210, USA; Department of Neurology, Cellular Biology and Anatomy, Louisiana State University Health Sciences Center, Shreveport, Louisiana 71130, USA; National Institute of Neuroscience, NCNP, 4-1-1 Ogawahigashi, Kodaira, Tokyo 187-8502, Japan

**Keywords:** Endothelial cells, monocytes, macrophages, hematopoietic growth factors, adhesion, stem cell factor, granulocyte colony-stimulating factor, chemokine receptor 5 perivascular macrophages

## Abstract

Monocytes are circulating macrophage precursors and are generated from bone marrow hematopoietic stem cells. In the adults, monocytes continuously replenish cerebral border-associated macrophages under a physiological condition. Monocytes also rapidly infiltrate into the brain in the settings of pathological conditions. The mechanisms of recruiting monocyte-derived macrophages into the brain under pathological conditions have been extensively studied. However, it remains unclear how monocytes enter the brain for renewal of border-associated macrophages under the physiological condition. Using both *in vitro* and *in vivo* approaches, this study reveals that the combination of two hematopoietic growth factors, stem cell factor (SCF) and granulocyte colony-stimulating factor (G-CSF), complementarily and synergistically enhances adhesion of monocytes to cerebral endothelial cells in a dose dependent manner. Cysteine-cysteine chemokine receptor 5 (CCR5) in brain endothelial cells, but not cell adhesion molecules mediating neuroinflammation-related infiltration of monocyte-derived macrophages, modulates the SCF+G-CSF-enhanced monocyte-endothelial cell adhesion. Blocking CCR5 or genetically deleting CCR5 reduces monocyte-endothelial cell adhesion induced by SCF+G-CSF. SCF+G-CSF-enhanced recruitment of bone marrow-derived monocytes/macrophages in cerebral perivascular space is also reduced in adult CCR5 knockout mice. This study demonstrates the contribution of SCF and G-CSF in regulating the entry of monocytes into the adult brain to replenish perivascular macrophages.

## 1. Introduction

The vascular endothelium, a monolayer of endothelial cells, forms the inner cellular wall of blood vessels and mediates many diverse biological processes including the maintenance of an anti-thrombotic surface between the circulating blood and tissue [1], regulation of vascular tone through the release of vasodilator and vasoconstrictive agents [2, 3], and selective separation of vascular luminal contents from the tissues through barrier formation [4]. The blood-brain barrier built on endothelial cells plays a vital role in maintaining the homeostasis of the brain microenvironment [5].

Macrophages in the brain consist of microglia in the parenchyma, border-associated macrophages, and monocyte-derived macrophages that enter the brain under pathological conditions [6]. Both the microglia and border-associated macrophages in the meningeal-choroid plexus-perivascular space are of embryonic origin, while the border-associated macrophages are renewed through monocyte recruitment under a physiological condition in the adult brain [6]. The mechanisms underlying the recruitment of monocyte-derived macrophages into the brain under pathological conditions have been extensively studied [5, 6]. However, it remains poorly understood how monocytes enter the adult brain to renew the border-associated macrophages under the physiological condition.

Stem cell factor (SCF, also termed master cell growth factor, kit ligand, and steel factor) and granulocyte colony-stimulating factor (G-CSF) are essential hematopoietic growth factors. SCF in combination with G-CSF shows synergistic effects in enhancing proliferation, differentiation, survival, and mobilization of hematopoietic stem cells [7–9]. SCF binds to its specific receptor, cKit, to regulate various biological functions such as hematopoiesis, gametogenesis, and melanogenesis [10–14]. G-CSF communicates with its receptor, GCSFR, to control proliferation, differentiation, and maturation of the precursor cells of granulocytes [15–17]. However, increasing evidence has revealed that SCF and G-CSF also have biological functions in the central nervous system. It has been shown that the combination treatment of SCF and G-CSF enhances neurogenesis and cerebral angiogenesis [18–21] and synergistically increases neurite outgrowth [22]. The expression of SCF and G-CSF receptors has been found on cerebral endothelial cells [23]. However, the exact biological function of SCF and G-CSF on cerebral endothelial cells has not been fully understood.

Monocytes expressing specific markers such as ionized calcium binding adaptor molecule 1 (Iba-1), CD115, and CD11b [24, 25] are a type of white blood cell and play multiple roles in the immune system, including replenishing resident macrophages and dendritic cells under normal states or pathological conditions [26, 27]. It is thought that monocyte recruitment in tissue may follow the general procedures of adhesion and trafficking, which includes rolling, adhesion, and transmigration of vascular endothelial cells [28, 29]. The adhesion molecules expressed on vascular endothelial cells are crucially involved in leukocyte infiltration during neuroinflammation under various disease conditions [5, 30]. It remains unclear, however, whether SCF and G-CSF could promote the expression of adhesion molecules or other molecules on cerebral vascular endothelial cells to recruit circulating monocytes into the brain and replace the border-associated macrophages under the physiological condition. Using both *in vitro* and *in vivo* approaches, the purpose of this study is to determine the effects and mechanism of SCF and G-CSF in regulating monocyte-endothelial cell adhension and the entry of monocytes into the adult brain under physiological conditions.

## 2. Results

### 2.1 Receptors for SCF and G-CSF are expressed on brain endothelial cells

To determine and validate the expressions of the receptors for SCF and G-CSF on brain endothelial cells, we performed immunofluorescence staining and flow cytometric analysis on adult mouse brain tissue and mouse brain-derived endothelial cells. The immunofluorescence staining in brain sections revealed that SCF receptor (Ckit) and G-CSF receptor (GCSFR) were co-localized with CD31^+^ endothelial cells in the cortex (Figure 1A and B). The flow cytometry described the mean fluorescence intensity (MFI) of Ckit as of 49.2 and G-CSFR as of 107 on the CD45^-^CD31^+^ gated primary endothelial cells isolated from adult mouse brain (Figure 1C and D). In addition, we also found that both Ckit and GCSFR were expressed on bEnd.3 cells (an endothelial cell line derived from mouse brain), which was confirmed by both immunofluorescence staining (Figure 1E and F) and flow cytometry (Figure 1G and H). These validated data demonstrate that the receptors for SCF and G-CSF are constitutively expressed on endothelial cells, suggesting that SCF and G-CSF might have important biological functions on the endothelial cells.

**Figure 1.**
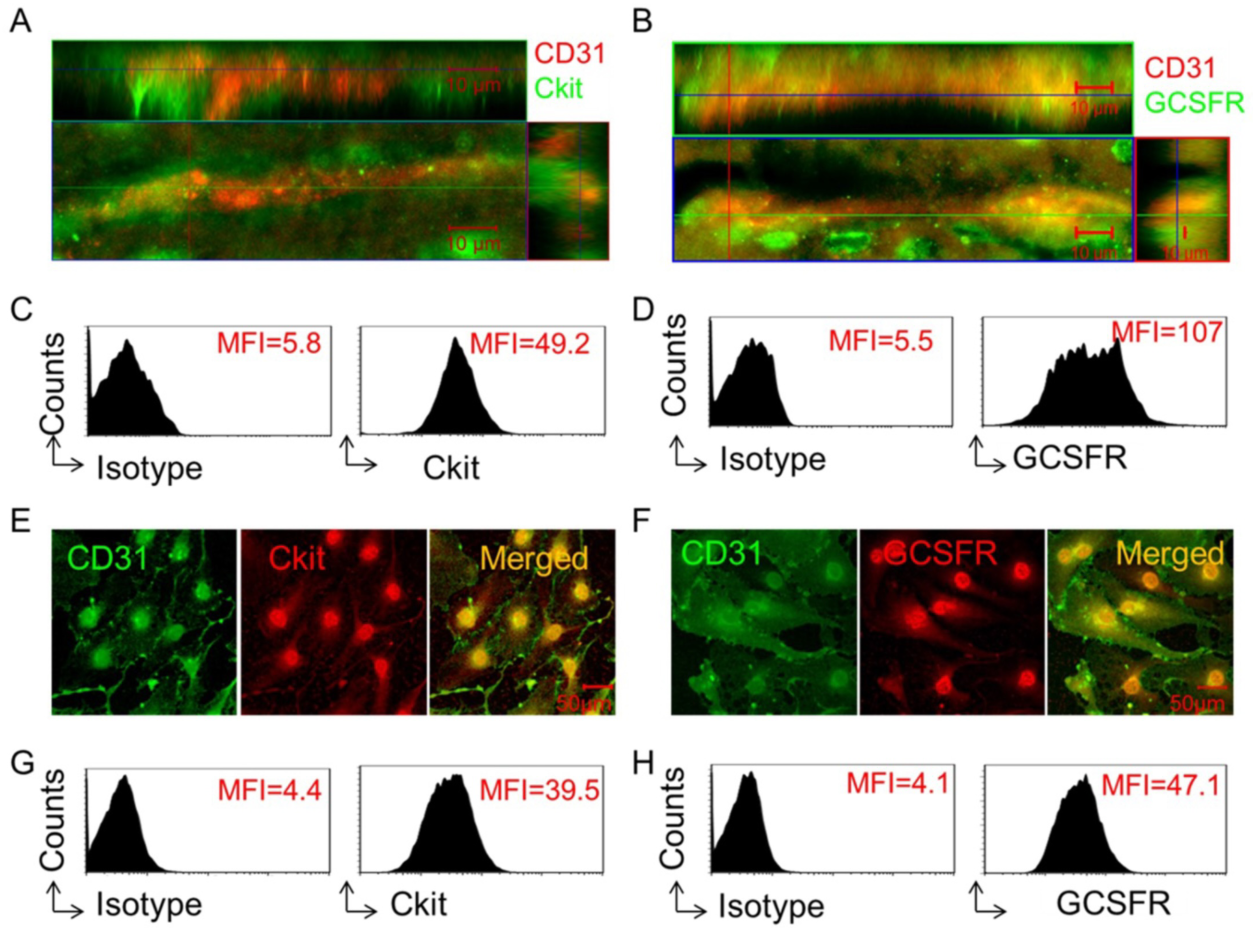
Receptors for SCF and G-CSF are expressed on brain endothelial cells. **(A, B)** Representative 3 dimensional (3D) confocal images. Capillaries (CD31^+^, red) in the cortex of adult mouse brain show co-expression with the receptors for SCF **(A)** (Ckit, green) and G-CSF **(B)** (GCSFR, green). Scale bar, 10μm. **(C, D)** Representative flow cytometry data. The expressions of Ckit **(C)** and GCSFR **(D)** on cerebral endothelial cells are detected by flow cytometry analysis of CD45 CD3V gated endothelial cells isolated from adult mouse brain. Mean fluorescence intensity (MFI) of Ckit (C) and GCSFR (D) expressing on cerebral endothelial cells is presented on flow cytometry histograms. Isotype: isotype control antibody. Repeated experiments, n = 4. **(E, F)** Representative confocal images of immunofluorescence staining. The expressions of Ckit **(E)** (red) and GCSFR **(F)** (red) are seen on bEnd.3 endothelial cells (CD31^+^, green). Scale bar, 50μm. Repeated experiments, n = 3. **(G, H)** Representative flow cytometry histograms showing the expressions of Ckit **(G)** and GCSFR (H) on bEnd.3 endothelial cells. MFI: Mean fluorescence intensity. Isotype: isotype control antibody. Repeated experiments, n = 5.

### 2.2 SCF in combination with G-CSF enhances monocyte adhesion to endothelial cells *in vitro*

As stated earlier, under the physiological condition, monocytes contribute to generating macrophages in adult brain [6, 31, 32]. However, it is not known how monocytes enter the adult brain. To address this question, we determined the effects of SCF and G-CSF in regulation of monocyte adhesion to endothelial cells.

To do so, we first examined whether SCF alone promotes the adhesion of monocytes to endothelial cells. To this end, we designed adhesion experiments using bEnd.3 cells and monocytes. Brain-derived endothelial cells (bEnd.3 cells) were pre-treated with SCF at different dosages for 16∼18h, and bone marrow-derived monocytes (Iba-1^+^ cells isolated from the bone marrow of Iba1-GFP mice) were then loaded on the top of the pre-treated bEnd.3 cells. After cells were incubated for 20 min, the adhesion assay was performed. We observed that significantly greater amounts of monocytes adhered to the endothelial cells pre-stimulated with SCF at 50ng/ml (*p* < 0.05) and 100ng/ml (*p* < 0.01) than those endothelial cells pre-treated with medium alone or a lower concentration of SCF (Figure 2A and D). These findings reveal that SCF enhances monocyte adhesion to endothelial cells.

**Figure 2.**
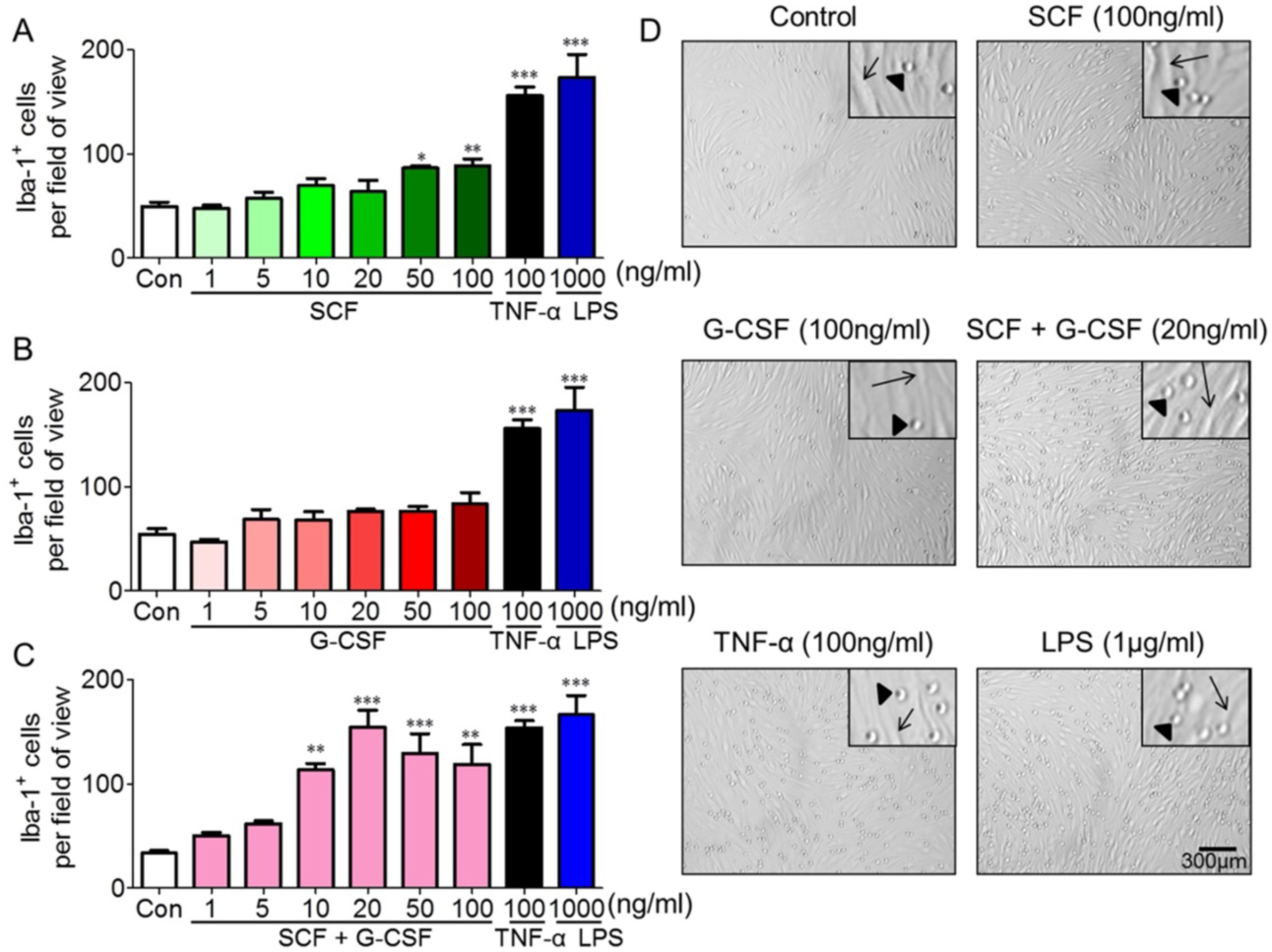
SCF in combination with G-CSF enhances lba-1^+^ monocyte adhesion to endothelial cells. **(A-C)** Monocyte-endothelial cell adhesion data, lba-1^+^ monocytes adhere to the bEnd.3 endothelial cells that are pre-treated with different doses of SCF alone (A), G-CSF alone **(B),** or SCF+GCSF **(C)** for 16-18 hours. Inflammatory mediators, TNF-α and LPS, serve as positive controls. Repeated by 3 independent experiments. Each experiment includes 3 samples (cell culture wells). Mean ± SEM. *\*p* < 0.05, **p < 0.01, ***p<0.001, vs. medium controls. One-way ANOVA followed by *post-hoc* Tukey’s test. (D) Representative bright-field images showing the adhesion of Iba-1^+^ monocytes on the bEnd.3 endothelial cells that are pre-treated with medium (control), SCF (lOOng/ml), GCSF (100ng/ml), SCF+G-CSF (20ng/ml), TNF-α (100ng/ml), and LPS (1μg/ml). The arrows show reference points illustrating spindle­shaped endothelial cells. The triangles indicate round lba-1^+^ monocytes. Scale bar, 300μm.

Next, we determined whether G-CSF alone has similar effects in enhancing endothelial cell/monocyte adhesion. Unexpectedly, G-CSF alone did not show significant changes in the adhesion of Iba-1^+^ monocytes to endothelial cells at all tested dosages (Figure 2B and 2D). By contrast, in the positive controls, significantly increased the numbers of monocytes adhering to bEnd.3 cells were observed following a 16-h pre-treatment with 100ng/ml TNF-α (*p* < 0.001) or 1µg/ml LPS (*p* < 0.001) as compared to the medium controls (Figure 2). The experiments were repeated four times, and similar results were obtained.

Further, we identified the efficacy of a combination of SCF and G-CSF (i.e. SCF+G-CSF) on endothelial cell/monocyte adhesion. Remarkably, SCF+G-CSF strongly enhanced Iba-1^+^ monocyte adhesion to bEnd.3 endothelial monolayers compared to medium controls (Figure 2C and D). The binding of monocytes to bEnd.3 endothelial cells reached its peak at the dose of 20ng/ml SCF and 20ng/ml G-CSF (*p* < 0.001). Although it still showed significant increases in monocyte/endothelial cell adhesion at the SCF and G-CSF doses of 50 ng/ml (p<0.001) and 100 ng/ml (p< 0.01), the levels of adhesion of monocytes to bEnd.3 cells slightly went down when the SCF+G-CSF dose increased (Figure 2C). These findings indicate that SCF+G-CSF enhances monocyte adhesion to endothelial cells in a dose dependent manner. Notably, SCF alone or G-CSF alone at the dose of 20ng/ml did not show increases in adhesion of Iba-1^+^ monocytes to endothelial cells (Figure 2A and B). To validate these findings, we used a flow cytometry approach to examine the differences between the adhesions of Iba-1-GFP^+^ monocytes to untreated/unstimulated bEnd.3 cells (control) and to the bEnd.3 cells pre-treated/pre-stimulated with SCF+G-CSF (20ng/ml), TNF-α (100ng/ml) or LPS (1µg/ml). As shown in the supplementary Figure1, Iba-1-GFP^+^ cells binding to SCF+G-CSF-pre-treated/pre-stimulated bEnd.3 cells were significantly increased compared to the untreated/unstimulated bEnd.3 cells (*p* < 0.01). Iba-1-GFP^+^ cells retained significantly higher adhesion ratio to the TNF-α or LPS pre-stimulated bEnd.3 cells than those untreated/unstimulated bEnd.3 cells (*p* < 0.001). These data strongly suggest that SCF in combination with G-CSF has a synergistic effect in promoting the adhesion function of endothelial cells.

### 2.3 SCF+G-CSF upregulates the expression of CCR5 but not adhesion molecules in the endothelial cells

Under inflammatory conditions, many adhesion molecules are expressed in the endothelial cells, these adhesion molecules are critically involved in rolling, adhesion, and migration of leukocytes across the vascular endothelial barrier [30, 33]. Here we used real-time qPCR and flow cytometry approaches to determine whether SCF+G-CSF increases the expression of adhesion molecules in the endothelial cells. We observed that gene expression levels of vascular cell adhesion molecule-1 (VCAM-1, CD106), intracellular adhesion molecule-1 (ICAM-1, CD54), P-selectin, and E-selectin were markedly up-regulated in bEnd.3 cells after 3h treatment with inflammatory mediators, TNF-α and LPS (*p* < 0.001) (Figure 3). However, SCF+G-CSF treatment did not significantly change the gene expressions of ICAM-1, VCAM-1, P-selectin, and E-selectin in the bEnd.3 cells (Figure 3). Flow cytometric analysis of ICAM-1 and VCAM-1 expression on the bEnd.3 cells was consistent with the observation from real-time qPCR (Figure 3 B and D). These findings suggest that the SCF+G-CSF-enhanced adhesion function of vascular endothelial cells is mediated through a pathway different from the inflammatory signaling.

**Figure 3.**
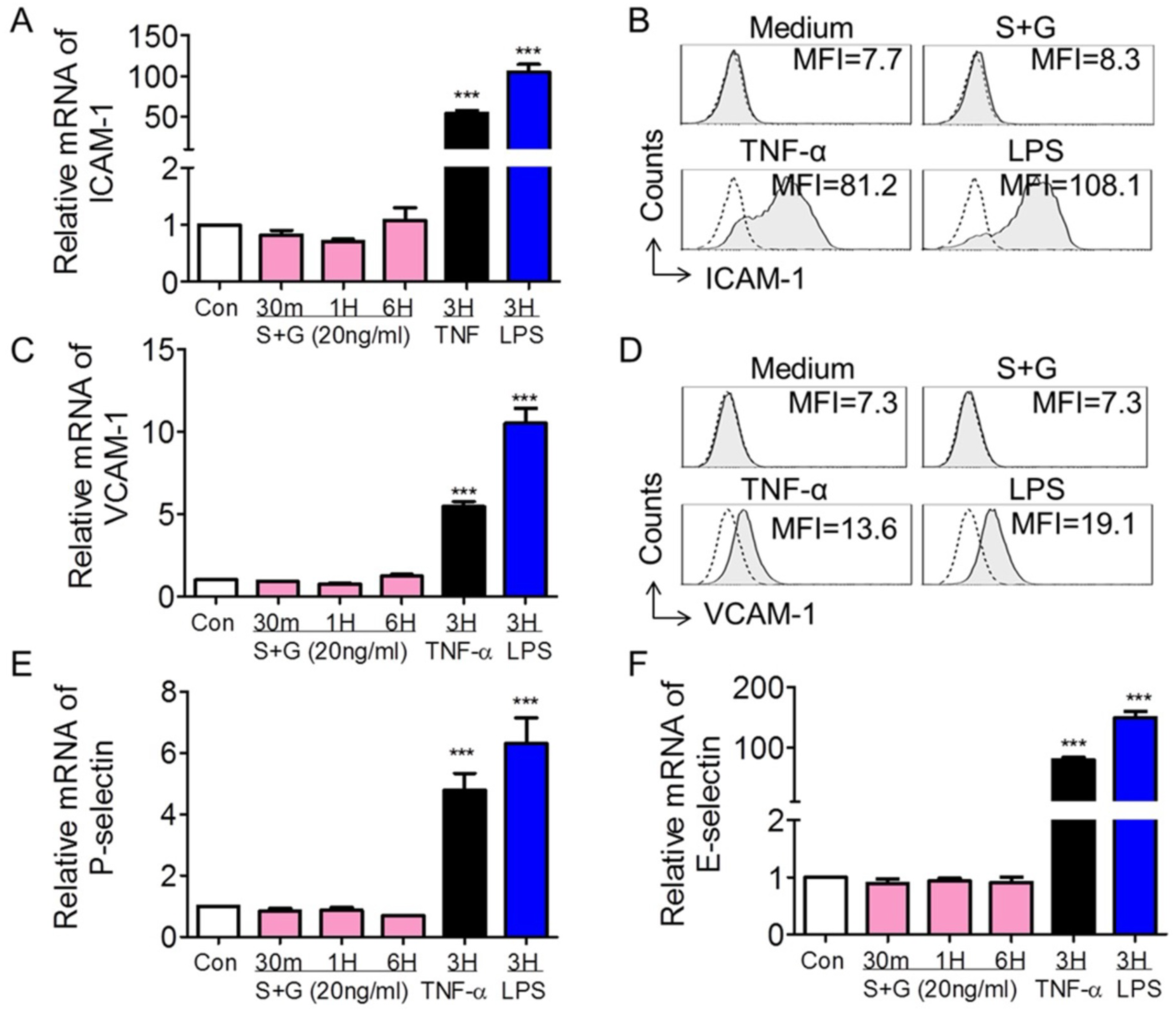
Expression of adhesion molecules in bEnd.3 endothelial cells. **(A, C, E, F)** Quantitative real-time PCR data. mRNA expressions of ICAM-1 **(A),** VCAM-1 **(C),** P-selectin **(E)** and E-selectin **(F)** in bEnd.3 endothelial cells that were cultured for different time periods (30min to 6 hours) in the presence of medium alone (con: control), SCF+G-CSF (20ng/ml), TNF-α (100ng/ml), and LPS (1μg/ml). Repeated by 3 independent experiments. Mean ± SEM. *\*\*\*p* < 0.001 vs. medium control. One-way ANOVA followed by *post-hoc* Tukey’s test. **(B, D)** Representative flow cytometry histograms showing the membrane expression levels of ICAM-1 **(B)** and VCAM-1 **(D)** on bEnd.3 cells with different treatments. MFI: Mean fluorescence intensity. Repeated experiments, n = 5. Dash lines indicate isotype IgG controls. Filled gray lines are specific staining for the indicated adhesion molecules.

Chemokines and chemokine receptors are structurally related proteins that regulate leukocyte adhesion and trans-endothelial migration both *in vitro* and *in vivo* [34, 35]. To further probe the possible mechanisms underlying the SCF+G-CSF-enhanced monocyte/endothelial cell adhesion, we examined the alterations of the chemokine receptors on bEnd.3 cells following SCF+G-CSF treatment using real-time qPCR. Our data showed that the mRNA levels of the chemokine receptors, CCR1, CCR2, CCR3, CCR4, CCR8, and CXCR4 were very low in bEnd.3 cells, and that 20ng/ml SCF+G-CSF treatment did not increase the mRNA expressions of these chemokine receptors (Supplementary Figure 2) with the exception of CCR5 when bEnd.3 cells were treated with 20ng/ml SCF+G-CSF. The mRNA expression of CCR5 in bEnd.3 cells was significantly increased by SCF+G-CSF treatment compared to medium controls (*p* < 0.001) (Figure 4A). This finding was further validated by flow cytometry. SCF+G-CSF treatment increased CCR5 expression on bEnd.3 cells (Figure 4B). Interestingly, the inflammatory mediators, TNF-α and LPS, significantly reduced CCR5 mRNA expression in bEnd.3 cells compared to medium controls (*p* < 0.001) (Figure 4A).

**Figure 4.**
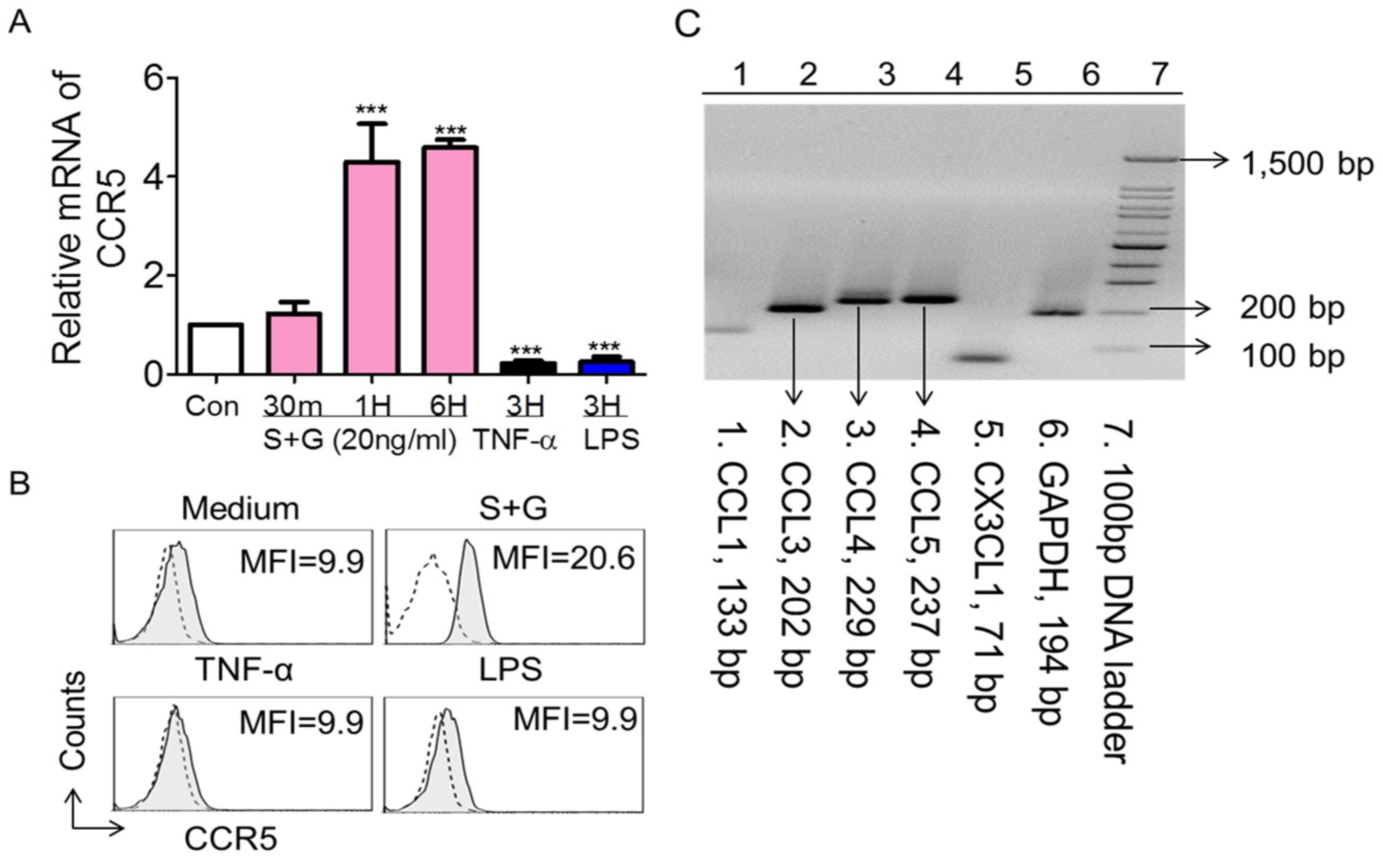
SCF+G-CSF increases CCR5 expression on bEnd.3 endothelial cells. **(A)** Quantitative real-time PCR data showing mRNA expression of CCR5 in bEnd.3 endothelial cells cultured for different time periods (30min to 6 hours) in the presence of medium alone (con: control), SCF+G-CSF (20ng/ml), TNF-α (100ng/ml), and LPS (1μg/ml). Mean ± SEM. \*\*\**p* < 0.001 *vs.* medium control. One-way ANOVA followed by *post-hoc* Tukey’s test. **(B)** Representative flow cytometry histograms showing the membrane expression of CCR5 on bEnd.3 endothelial cells. MFI: Mean fluorescence intensity. Repeated experiments, n = 5. Dash lines indicate isotype IgG controls. Filled gray lines are specific staining for CCR5. **(C)** PCR-electrophoresis showing mRNA expressions of CCL1 (133 bp), CCL3 (202 bp), CCL4 (229 bp), CCL5 (237 bp), and CX3CL1 (71 bp) in lba-1^+^ monocytes.

### 2.4 SCF+G-CSF enhances monocyte adhesion to endothelial cells through CCR5 *in vitro*

CCR5 binds to several chemokine ligands including CCL3, CCL4, and CCL5 [36–38]. In studying the cell-cell adhesion, it is important to detect whether Iba-1^+^ monocytes express the ligands of CCR5 when CCR5 receptors are increased on the endothelial cells by SCF+G-CSF. Our data revealed that Iba-1^+^ monocytes constitutively express CCL3, CCL4, and CCL5 (Figure 4C). These data suggest that the CCR5-related chemokine-receptor pathway could be an important mechanism underlying the adhesion of monocytes to SCF+G-CSF-treated endothelial cells.

We then sought to determine the role of endothelial CCR5 in SCF+G-CSF-enhanced monocyte-endothelial adhesion. To block CCR5, after pre-treatment of bEnd.3 monolayers with SCF+G-CSF (20µg/ml), anti-CCR5 antibody (50µg/ml) or its isotype IgG2a control (50µg/ml) was added to the washed bEnd.3 cells. Antibodies were washed out with pre-warmed medium, and then FACS-sorted Iba-1-GFP^+^ monocytes were added to the bEnd.3 monolayers. Remarkably, the SCF+G-CSF-increased monocyte/endothelial cell adhesion was completely blocked by anti-CCR5. The control antibody against IgG2a did not show an influence on adhesion of monocytes to SCF+G-CSF-treated bEnd.3 cells (Figure 5). The anti-CCR5 antibody did not block the adhesion of Iba-1^+^ monocytes to the inflammatory cytokine TNF-α-stimulated bEnd.3 cells. These data strongly suggest that endothelial CCR5 controls the adhesion of monocytes to the SCF+G-CSF-pre-treated endothelial cells, which is completely different from an inflammatory signaling-mediated pathway for monocyte-endothelial cell adhesion.

**Figure 5.**
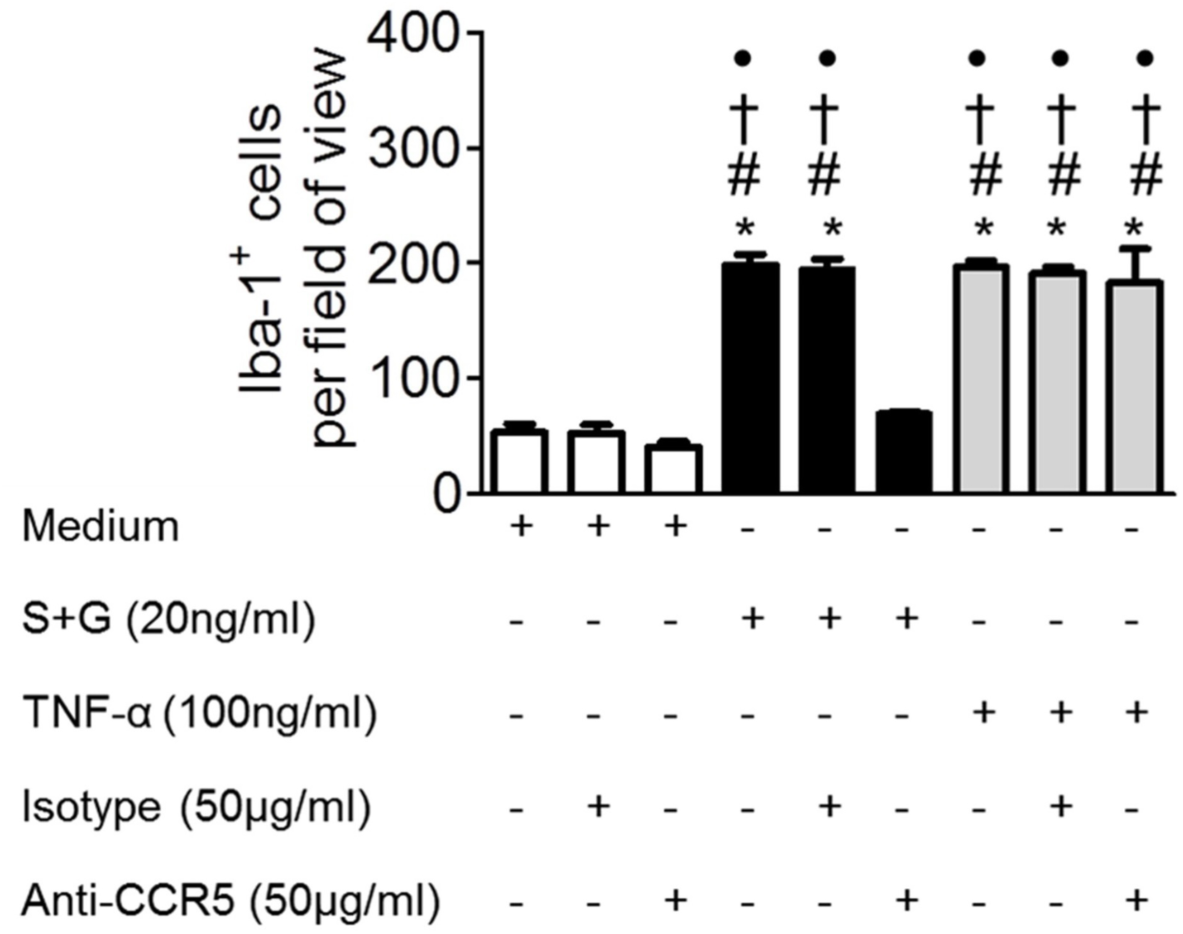
Anti-CCR5 antibody blocks SCF+G-CSF-enhanced monocyte adhesion to endothelial cells. The monolayers of mouse brain-derived endothelial cells (i.e. bEnd.3 cells) were incubated in the presence of medium alone, SCF+G-CSF (20ng/ml), and TNF-α (100ng/ml) for 16-18 hours. After washing, the bEnd.3 cells were pre­treated with anti-CCR5 antibody or Isotype IgG control antibody before adding lba-1^+^ monocytes to the bEnd.3 cell monolayers. Mean ± SEM. N = 7. \**p* < 0.001 vs. medium control, #*p* < 0.001 *vs.* medium with Isotype control, †*p* < 0.001 *vs.* medium with anti-CCR5, and \**p* < 0.001 *vs.* S+G with anti-CCR5. One-way AN OVA followed by *post-hoc* Tukey’s test.

### 2.5 SCF+G-CSF enhances bone marrow-derived cells adhering to cerebral endothelial cells through CCR5 *in vivo*

To verify that SCF+G-CSF treatment increases blood cell adhesion to endothelial cells under a physiological condition *in vivo*, we created UBC-GFP^+^ bone marrow chimeric CCR5^-/-^ and WT mice (i.e. UBC-GFP-CCR5^-/-^ mice and UBC-GFP-WT mice). In these mice, we examined the adhesion of bone marrow-derived cells (GFP positive) on cerebral endothelial cells (CD31 positive) after SCF+G-CSF treatment (experimental flow chart, Figure 6A). We observed that SCF+G-CSF treatment increased bone marrow-derived GFP^+^ cell adhesion to CD31^+^ endothelial cells in the cerebral cortex of both UBC-GFP-WT mice and UBC-GFP-CCR5^-/-^ mice compared to the vehicle control groups (*p* < 0.001). However, the number of GFP^+^ cells adhering to endothelial cells was significantly less in SCF+G-CSF-treated UBC-GFP-CCR5^-/-^ mice than in the UBC-GFP-WT mice treated with SCF+G-CSF (*p* < 0.001) (Figure 6). These findings suggest that the SCF+G-CSF-enhanced adhesion of bone marrow-derived blood cells to brain endothelial cells is, at least partially, mediated through CCR5.

**Figure 6.**
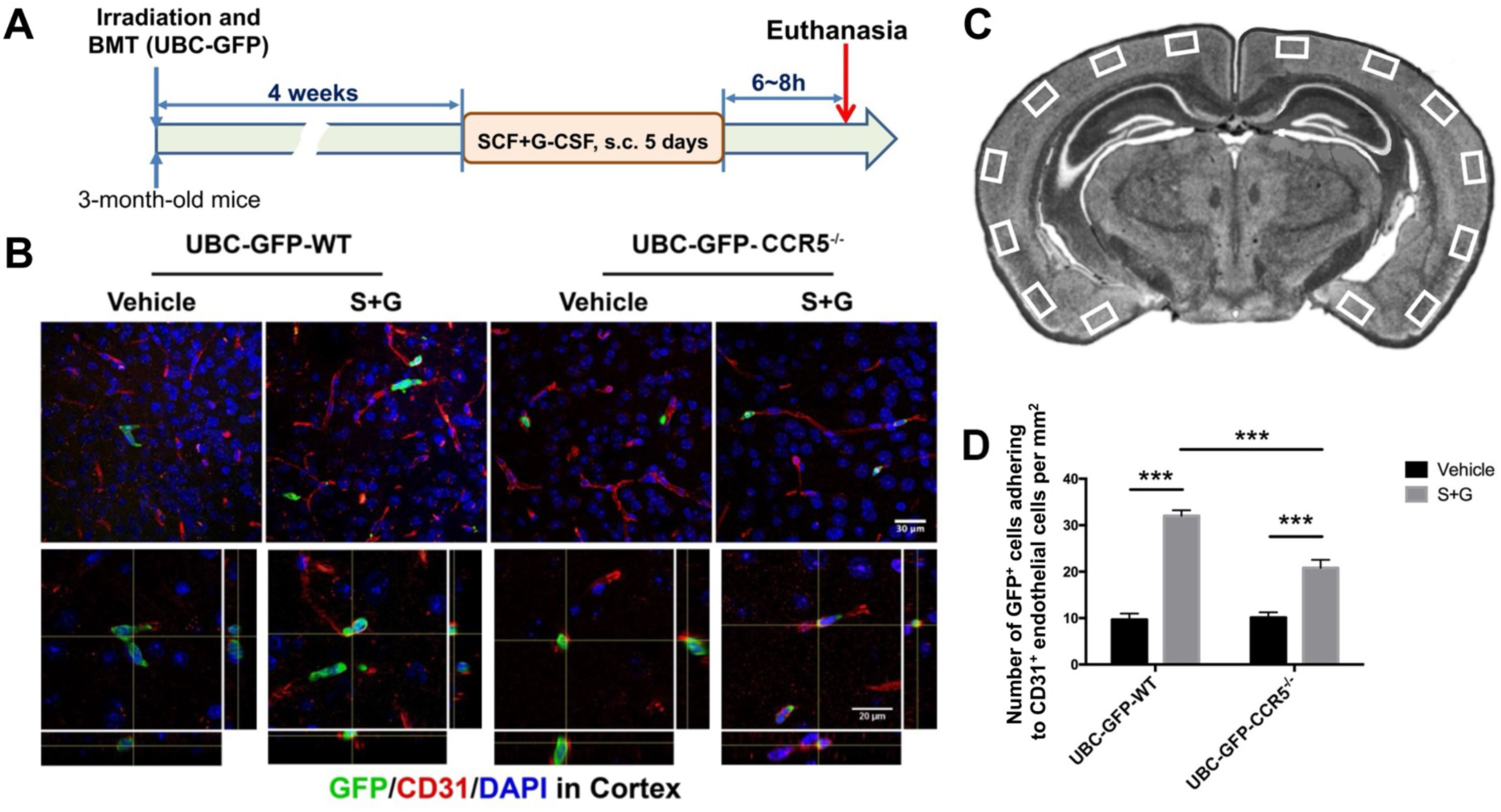
SCF+G-CSF increases adhesion of bone marrow-derived cells to cerebral endothelial cells partially through CCR5 in adult mice. **(A)** Experimental flow chart. BMT: bone marrow transplantation. Bone marrow donor: UBC-GFP mice. **(B)** Representative confocal images showing immunofluorescence staining of GFP positive bone marrow-derived cells (green), and CD31 positive endothelial cells (red) in the cortex of UBC-GFP bone marrow-transplanted vehicle-control WT mice, vehicle-control CCR5^-/-^ mice, SCF+G-CSF-treated WT mice, and SCF+G-CSF-treated CCR5^-/-^ mice. Z-stack images of 15 layers with 1μm intervals. DAPI (blue): nuclear counterstaining. Upper 4 images: Projection views of z-stack images (Scale bar, 30 μm). Lower 4 images: Orthogonal views of z-stack images showing 3D visualizations of the colocalizations of bone marrow-derived cells (GFP, green) with endothelial cells (CD31, red). Scale bar, 20 μm. **(C)** A diagram that shows the selected regions in the cortex for confocal imaging. **(D)** Quantification data that show the number of GFP^+^ cells that adhere to CD31^+^ endothelial cells per mm^2^ in experimental groups. Mean ± SEM. N=5, *\*\*\*p* < 0.001. One-way ANOVA followed by *post-hoc* Tukey’s test.

### 2.6 CCR5 mediates bone marrow-derived monocyte transmigration into the brain surrounding the blood vessels

To identify whether CCR5 modulates bone marrow-derived monocyte transmigration into the brain by SCF+G-CSF treatment, we quantified the number of GFP^+^/Iba1^+^ cells surrounding the CD31^+^ endothelial cells in the cortex (surrounding: within 10 μm from vessels). In the vehicle control mice, the number of GFP^+^/Iba1^+^ cells surrounding the blood vessels in the cerebral cortex was significantly reduced in CCR5 knockout mice compared to WT mice (*p* < 0.05) (Figure 7). Similarly, in the SCF+G-CSF-treated mice, the number of GFP^+^/Iba1^+^ cells surrounding the blood vessels in the cortex was significantly decreased in CCR5 knockout mice compared to WT mice (*p* < 0.05) (Figure 7). These data suggest that CCR5 is involved in the transmigration of Iba1^+^ bone marrow-derived monocytes into the brain, and that CCR5 is required for SCF+G-CSF-enhanced transmigration of blood monocytes into the brain. Interestingly, GFP^+^/Iba1^+^ cells were found only around but not inside the lumen of CD31^+^ brain vessels, indicating that monocytes may differentiate into Iba1^+^ macrophages right after transmigrating from the blood vessels into the brain.

**Figure 7.**
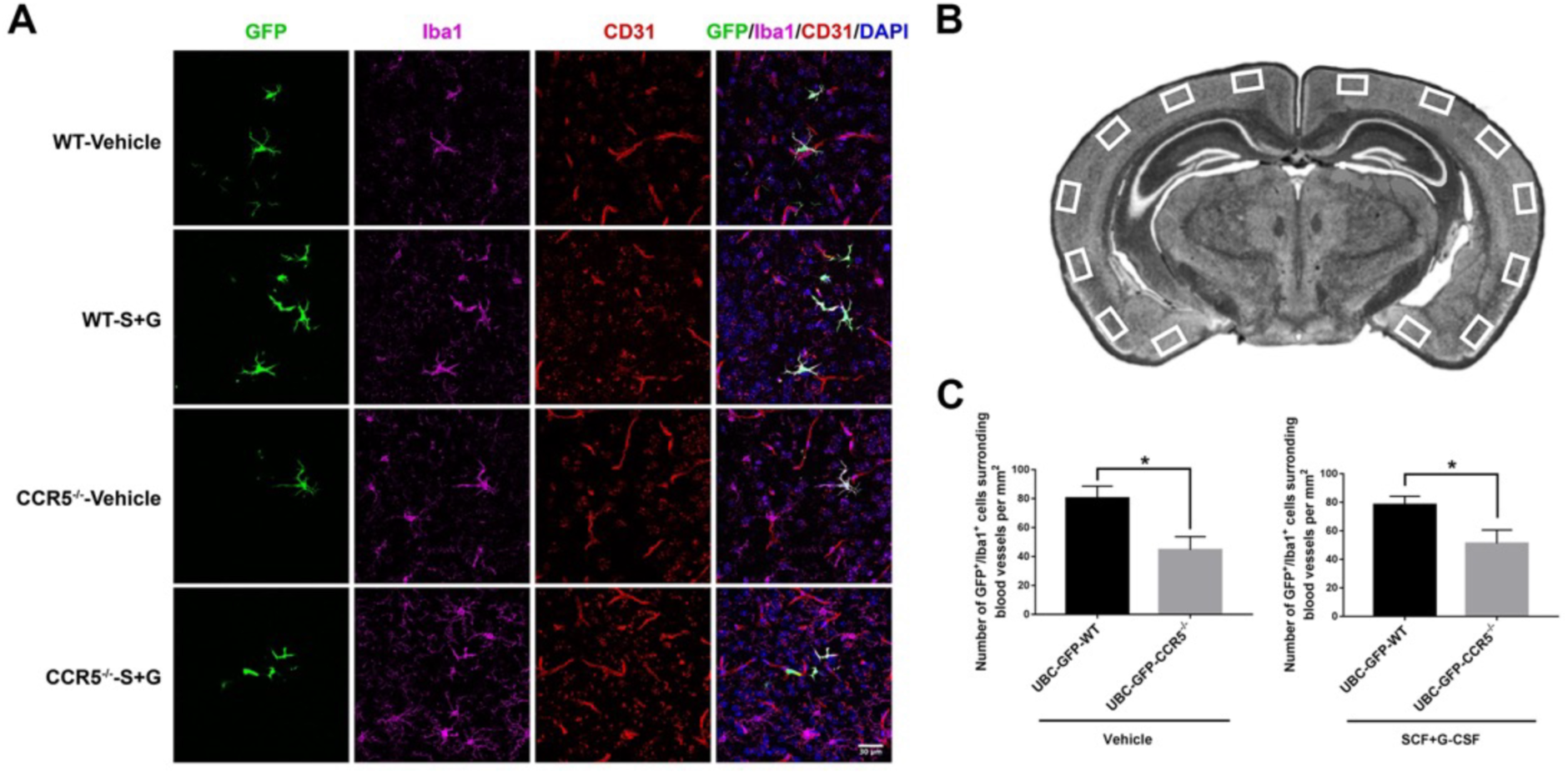
CCR5 mediates bone marrow-derived monocyte transmigration into the adult brain. **(A)** Representative projection views of z-stack confocal images that show the location of bone marrow-derived monocytes (GFP^+^/lba1^+^ cells) next to the blood vessels (CD31^+^) in cerebral cortex of adult mice that received UBC-GFP bone marrow transplantation in the groups of vehicle-control WT mice, vehicle-control CCR5^-/-^ mice, SCF+G-CSF-treated WT mice, and SCF+G-CSF-treated CCR5^-/-^ mice. DAPI (blue): nuclear counterstaining. Scale bar, 30μm. Z-stack images of 15 layers with 1 μm intervals. **(B)** A diagram that shows the selected regions in the cortex for confocal imaging. **(C)** Quantification data. The number of GFP^+^/lba1^+^ bone marrow-derived monocytes/macrophages surrounding CD31^+^ blood vessels (within 10μm from the blood vessels) in the cerebral cortex is reduced in both vehicle-control CCR5^-/-^ mice and SCF+G-CSF-treated CCR5^-/-^ mice. Mean ± SEM. N=5. *\*p* < 0.05, Unpaired t-test.

## 3. Discussion

In the present study, we have, for the first time, identified that the combination of two hematopoietic growth factors, SCF and G-CSF, can enhance the replenishment of perivascular macrophages in adult mouse brain via CCR5 under the physiological condition. The findings of our *in vitro* experiments reveal that SCF in combination with G-CSF (SCF+G-CSF) complementarily and synergistically promotes the adhesion function of brain endothelial cells to bind with bone marrow-derived monocytes. The SCF+G-CSF-enhanced adhesion function of brain endothelial cells is mediated through the chemokine receptor 5 (CCR5), but not cell adhesion molecules, which is completely different from an inflammatory signaling-triggered cell adhesion molecule pathway for monocyte-endothelial cell adhesion. The findings of our *in vivo* experiment further confirm the role of CCR5 in modulating SCF+G-CSF-enhanced renewal of perivascular macrophages in the adult brain by bone marrow-derived monocytes in the physiological state.

SCF and G-CSF act on their target cells through specific membrane receptors to regulate many cell functions including promoting neurite outgrowth [22, 39] and inhibiting apoptosis [40]. In the present study, we have found that the receptors for SCF and G-CSF are expressed in endothelial cells of the adult mouse brain and in the brain-derived endothelial cell line. This observation is in line with the previous report showing that SCF receptor, cKit, and GCSF receptor, GCSFR, are expressed in cerebral endothelial cells [23]. These findings implicate the ability of SCF and G-CSF to modulate endothelial cell functions.

Using the well-established cell adhesion model *in vitro*, our novel finding reveals that SCF+G-CSF shows a synergistic effect on monocyte-endothelial cell adhesion greater than either of them alone. The best dose to increase monocyte-endothelial cell adhesion reached a peak at 20ng/ml of SCF+G-CSF *in vitro*, and the number of adherent monocytes was not further increased by dose increase. It is worth noting that SCF in combination with G-CSF also shows synergistic effects in enhancing the proliferation, differentiation, survival, and mobilization of hematopoietic stem cells [7–9] and in promoting neurite outgrowth [22]. The mechanism underlying the synergistic effects of SCF+G-CSF in the regulation of cell functions is not clear. One possibility could be that the proper doses of SCF+G-CSF in increasing monocyte-endothelial cell binding may robustly activate their receptor-mediated cell signaling because either SCF-Ckit or G-CSF-GCSFR interaction leads to multiple signaling cascades including the RAS/ERK, PI3-K/AKT, Src Kinase, JAK/STAT and MEK/ERK pathways [22, 41–43]. Both SCF and G-CSF share some similar signaling pathways but each of them could have relatively independent signaling pathways. Our data show that G-CSF alone does not affect adhesion of monocytes to endothelial cells when the endothelial cells lack the needed signaling triggered by SCF. The precise mechanism by which this occurs remains unclear. It has been shown that SCF+G-CSF synergistically increases myeloid cell proliferation through complementary signaling pathways [7]. G-CSF, but not SCF, induces the tyrosine phosphorylation of STAT1 and STAT3 signaling. However, SCF induces phosphorylation of STAT3 on serlne727, which is required for maximal and complete STAT3 transcriptional activity when SCF is combined with G-CSF [7]. The underlying molecular mechanisms of the SCF+G-CSF-induced synergistic effects in enhancing endothelial cell adhesion function when binding with monocytes, however, need further investigation.

It is widely known that inflammatory signals initiate the expressions of the adhesion molecules on the endothelial cell surface (such as ICAMs, VCAMs, P-selectin and E-selectin) [30, 34, 44]. Adhesion molecules are responsible for the leukocyte-endothelial cell crosstalk and mediate the entry of leukocytes into the tissue under pathological conditions [45–49]. How leukocyte-endothelial cell adhesion unfolds during the initial phase is not entirely clear, but leukocyte recruitment is undoubtedly of fundamental importance and continues to be a predominant feature during inflammation [50]. Remarkably, our study demonstrates that SCF+G-CSF fails to induce the changes of inflammation-related cell adhesion molecules, ICAM, VCAM, P-selectin or E-selectin although SCF+G-CSF leads to robust increases in monocyte-endothelial cell adhesion. These data open the possibility that the molecules mediating the adhesion of monocytes to endothelial cells by SCF+G-CSF are completely different from inflammation-induced cell adhesion. As confirmed by the findings of our *in vivo* experiment, SCF+G-CSF treatment increases cerebral endothelial cell adhesion to bone marrow-derived cells in heathy adult wild-type mice, supporting the role of SCF+G-CSF in enhancing monocyte-endothelial cell adhesion under a physiological condition.

It has been demonstrated that chemokines and chemokine receptors regulate leukocyte adhesion and trans-endothelium migration both *in vitro* and *in vivo* [51–53]. Activation signals of chemokines and chemokine receptors initiate firm adhesion of leukocytes to endothelial cells by rapidly upregulating integrin affinity [54, 55]. In the present study, we screened chemokine receptors expressed by endothelial cells with SCF+G-CSF stimulation. Our data show that CCR5 is pivotal in controlling the monocyte-endothelial cell adhesion induced by SCF+G-CSF. It has been shown that CCR5 is expressed in different types of cells, including endothelial cells [56]. The natural chemokine ligands that interact with CCR5 are CCL5 (also known as RANTES), macrophage inflammatory protein (MIP)-1α and MIP-1β (also known as CCL3 and CCL4), and CCL3L1 [36–38, 57]. In the present study, the expressions of these CCR5 chemokine ligands are detectable on bone marrow-derived Iba-1^+^ monocytes, suggesting the possibility of monocyte-endothelial cell interaction via CCR5-related ligand-receptor binding. Our data also reveal that the expression of CCR5 on endothelial cells is upregulated by treatment with SCF+G-CSF *in vitro*. Blocking CCR5 on endothelial cells eliminates the SCF+G-CSF-induced binding of monocytes to endothelial cells, demonstrating that CCR5 plays an important role in the SCF+G-CSF-enhanced adhesion of monocytes to endothelial cells. These findings are further validated by our *in vivo* study. CCR5 knockout mice which received bone marrow transplantation from UBC-GFP mice exhibited decreased adhesion of bone marrow-derived GFP^+^ cells to endothelial cells induced by SCF+G-CSF treatment compared with WT mice. However, CCR5 knockout only partially blocks the SCF+G-CSF-induced adhesion of bone marrow-derived cells to endothelial cells. There might be other underlying mechanisms by which SCF+G-CSF treatment enhances adhesion of bone marrow-derived blood cells to the cerebral endothelial cells. In addition to the interaction of CCR5 and its ligands, it is possible that other ligand-receptor interactions may also participate in SCF+G-CSF-enhanced monocyte adhesion to endothelial cells, such as the EphrinB2/EphA4 interaction [58]. It is worth mentioning that in our *in vitro* experiments blocking CCR5 fails to reduce monocyte-endothelial cell adhesion triggered by TNF-α or LPS (inflammatory mediators), indicating that inflammation-related monocyte-endothelial cell adhesion does not require CCR5, which is different from SCF+G-CSF-enhanced monocyte-endothelial cell adhesion.

Interestingly, in our *in vivo* study, bone marrow-derived Iba1^+^ cells were only observed surrounding but not in the lumen of brain blood vessels, indicating that bone marrow-derived monocytes express Iba1 or the bone marrow-derived monocytes differentiate into macrophages only after transmigrating from the blood vessels into the brain.

The mechanisms by which monocytes adhere to cerebral endothelial cells and transmigrate into to the perivascular space could be under the coordinated control of a wide range of signal pathways, many of which are only just beginning to be understood. The findings presented here not only demonstrate that SCF+G-CSF plays a pivotal role on monocyte-endothelium adhesion and transmigration, but also further substantiate that CCR5 is involved in mediating the SCF+G-CSF-enhanced monocyte-endothelium adhesion and transmigration. These findings shed new light on the understanding of the biological function of SCF+G-CSF in acting on the endothelial cells, and a regulatory role of SCF+G-CSF in replenishing perivascular macrophages from monocytes in the adult brain under the physiological condition.

## 4. Material and methods

### 4.1. Animals

All experiments were carried out in accordance with protocols approved by Institutional Animal Care and Use Committee and in keeping with the National Institute of Health’s Guide for the Care and Use of Laboratory Animals.

For the *in vitro* study, transgenic mice carrying Iba1-*enhanced green fluorescent protein* (GFP) with a C57BL/6 background, originally developed by Dr. Kohsaka [59], were used. The original breeding pairs were kindly provided by Dr. Kohsaka. C57BL/6 mice (Jackson Laboratory, Bar Harbor, ME, USA) served as wild-type (WT) controls. Male mice at the age of 8-10 weeks were used for experiments.

For the *in vivo* study, transgenic mice ubiquitously expressing enhanced GFP under the control of the human ubiqutin C promoter (UBC-GFP) with a C57BL/6 background, CCR5 knock out (CCR5^-/-^) mice (C57BL/6 genetic background), and C57BL/6 mice were purchased from Jackson Laboratory (Bar Harbor, ME, USA). Male mice were used for experiments.

Mice were housed under a 12-h light/dark cycle with *ad libitum* access to food and water.

### 4.2. Culture of bEnd.3 endothelial cells

The murine brain endothelial cell line bEnd.3 (ATCC, CRL-2299) was cultured in DMEM medium (ATCC, Manassas, VA, USA) with 10% fetal bovine serum (FBS; Atlanta Biologicals, Atlanta, GA, USA), 100U/ml penicillin (Invitrogen), and 100µg/ml streptomycin (Invitrogen) at 37°C under a humidified atmosphere of 5% CO2. Confluent monolayers were passed by adding 0.25% trypsin-EDTA (Cellgro, Manassas, VA, USA) into 24 well cell culture plates.

The bEnd.3 cells at passages 20 to 30 were used for adhesion assays or mRNA analysis when they formed confluent endothelial cell monolayers.

### 4.3. Isolation and culture of brain endothelial cells

C57BL/6 mice were deeply anesthetized and transcardially perfused with 30 ml of phosphate-buffered saline (PBS) to remove blood cells. The forebrain was removed and suspended in RPMI-1640 medium (ThermoFisher Scientific, Liverpool, NY, USA). The suspension was digested with type I collagenase (1mg/ml, Worthington, NJ, USA) and DNase I (50 µg/ml, Roche Diagnostics, Indianapolis, IN, USA) at 37°C for 45 min in a shaker set at 180 revolutions per minute. Brain endothelial cells were isolated using 37– 70% Percoll (GE Healthcare, Chicago, IL, USA) density gradient centrifugation according to a method described previously [60]. The endothelial cells were obtained from the interface, washed twice with DMEM medium, and resuspended in DMEM medium containing 10% FBS for phenotyping.

### 4.4. Isolation of Iba-1 positive monocytes from bone marrow cells

Iba1^+^ monocytes were obtained from the bone marrow of Iba1-GFP mice. Iba1-GFP mice were used for isolation of Iba1^+^ monocytes in this study because Iba1 is expressed in blood monocytes [61, 62]. The Iba1-GFP mice were anesthetized and euthanized by cervical dislocation. The femur and tibia bones were removed, cleaned of all connective tissue, and placed on ice in complete bone marrow medium (CBMM). CBMM consisted of DMEM medium supplemented with 10% horse serum (HS; Hyclone Laboratories, Logan, UT, USA). The ends of each femur and tibia bone were clipped to expose the marrow. Syringes with 21-gauge needles were used to flush out the bone marrow cells from one end of the bone until it turned completely white. The cells were suspended in CBMM and centrifuged at 1500 rpm for 5 minutes at 4°C. The cells were then resuspended in red cell lysis buffer (eBioscience, San Diego, CA, USA) and incubated at room temperature for 5 min. The cells were filtered through a 70µm nylon mesh strainer (BD, Franklin Lakes, NJ, USA) and then washed three times with CBMM. The Iba-1^+^ monocytes were further sorted by Fluorescence Activated Cell Sorting (FACS, BD FACSVantage Diva, USA). Post-sorting reanalysis showed > 95% purity of the Iba-1-GFP^+^ population.

### 4.5. Analysis of cells by flow cytometry

Anti-mouse antibodies used for flow cytometry included: rat anti-CD31 (MEC 13.3, BD Pharmingen, San Diego, CA, USA), rat anti-CD45-APC (30F11, eBioscience, San Diego, CA, USA), rabbit anti-mouse cKit (sc-168, Santa Cruz Biotechnology, Dallas, TX, USA), rabbit anti-mouse GCSFR (sc-694, Santa Cruz Biotechnology, USA), rabbit IgG (sc-3888, Santa Cruz Biotechnology, USA), anti-CCR5-PE (HM-CCR5 (7A4), eBioscience, USA), anti-CD106-FITC (VCAM-1, 429, eBioscience, USA), and anti-CD54-FITC (ICAM-1, YN1/1.7.4, eBioscience, USA). The brain single cell suspension was centrifuged, and the supernatant was removed. The cell pellets were fixed in cold 4% paraformaldehyde (PFA) (Polysciences, Inc. Warrington, PA, USA) at room temperature for 20 min and washed with staining medium (PBS containing 0.1% NaN3 and 2% FCS). The cells were incubated with primary antibodies (combinations of anti-CD31 and anti-cKit, anti-CD31 and anti-GCSFR, anti-CD31 and rabbit IgG) for 1 hour and washed twice with staining medium. The cells were then incubated with Cy2-anti-Rat (1:200, Jackson ImmunoResearch Laboratories, West Grove, PA, USA) and Dylight-549-conjugated anti-Rabbit (diluted 1:400, Jackson ImmunoResearch Laboratories) for 30 min. The cells were washed again and incubated with anti-CD45-APC for an additional 30 min. An extra cell wash was performed before detection by flow cytometry (FACSCalibur, BD science). For bEnd.3 cells, the cultured cells were rinsed with 0.25% trypsin-EDTA to get a single cell suspension and processed following the same staining procedures as stated above. Data were analyzed using FlowJo software (TreeStar, Ashland, OR, USA). Brain endothelial cells were gated on CD45^-^/CD31^+^ population and mean fluorescence intensity (MFI) was presented relative to appropriate isotype controls.

### 4.6. Adhesion assay

The adhesion experiments were performed as previously described [63]. bEnd.3 monolayers were incubated with stem cell factor (SCF) (PeproTech, Rocky Hill, NJ, USA) and/or granulocyte colony-stimulating factor (G-CSF) (Amgen, Thousand Oaks, LA, USA) for 16 to 18 hours at the concentration indicated. TNF-α (100ng/ml) (Life Technologies, Grand Island, NY, USA) and LPS (1µg/ml) (Sigma, St. Louis, MO, USA) served as positive controls. Stimulated bEnd.3 cells were washed three times with pre-warmed 10% fetal calf serum (FCS) in DMEM. Iba1^+^ monocytes (500,000 cells in a volume of 200ul per well) were then added on the top of the bEnd.3 cells. In some experiments, stimulated bEnd.3 cells were incubated with anti-mouse-CCR5 (10µg/ml) (HM-CCR5 (7A4), eBioscience, USA) or Armenian Hamster IgG isotype control antibody (eBio299Arm, eBioscience, USA) for 30 min at 37°C and washed before Iba1^+^ monocytes were added to the bEnd.3 cells. The cell culture plates were incubated at room temperature on a horizontal shaker (70rpm) for 20 min. The plates were washed twice to remove unbound or loosely bound cells and fixed in 4% PFA in PBS for 20 min. The determination of binding was made by counting the adherent GFP^+^ cells in four randomly chosen microscope fields (X20) in each well. The cells of four fields were averaged for data analysis. Data collection was blinded to the experimental treatments.

### 4.7. Real-time quantitative polymerase chain reaction

Total RNA was extracted from cultured cells using a RNeasy Plus Mini Kit (Qiagen, Hilden, Germany). First strand complementary DNA (cDNA) was prepared from an RNA template (1 µg) using the High-Capacity RNA-to-cDNA kit (Applied Biosystems, Waltham, MA, USA). Reverse transcription was performed at 37°C for 60 min and then at 95°C for 5 min following manufacturer’s instructions. Using the SsoAdvanced SYBR Green Supermix (Bio-Rad, Hercules, CA, USA), real-time quantitative polymerase chain reaction (RT-qPCR) amplification was performed by enzyme activation at 95°C for 30s and denaturation at 95°C for 5s, and annealing and extension at 58°C for 1min. Forty cycles of this process were repeated, and then melting curve analyses were performed for each reaction to confirm single amplified products between 65-95°C in 0.2°C increments on the Bio-Rad CFX96 system (Bio-Rad, USA). Relative quantification of mRNA of target genes across several time points was determined by taking the ratio of CT values divided by the CT values of glyceraldehyde-3-phosphate dehydrogenase (GAPDH) acquired from the corresponding time. All the primers were ordered from Integrated DNA Technologies. Primer sequences are listed in Table 1.

**Table 1:**
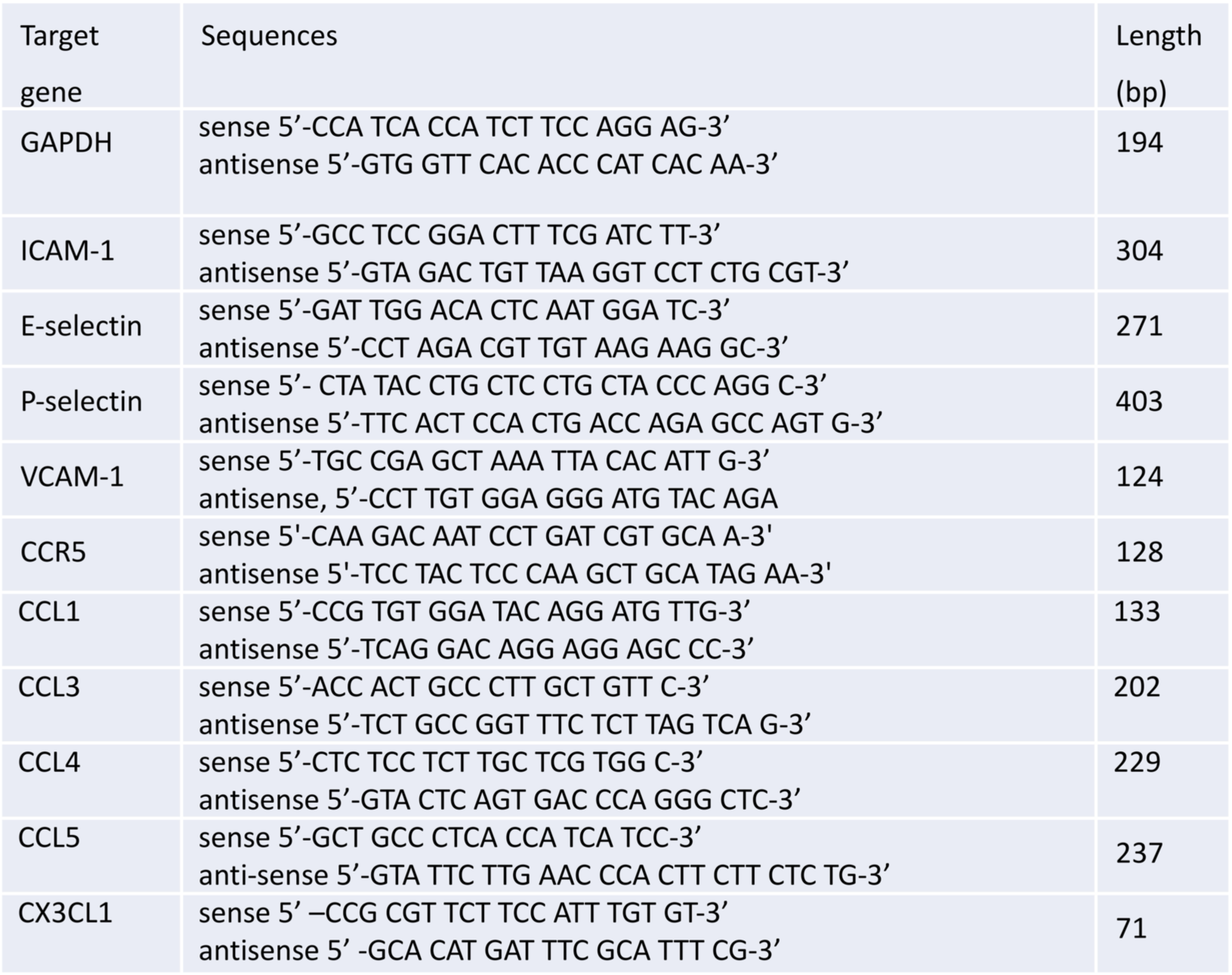
Primers for PCR.

### 4.8. Experimental Design of *In Vivo* Study

Mice were divided into four experimental groups at 3 months of age: a WT-vehicle control group (n = 5), a WT-SCF+G-CSF group (n = 5), a CCR5^-/-^-vehicle control group (n = 5), and a CCR5^-/-^-SCF+G-CSF group (n = 5). After mice received a lethal dose of whole-body irradiation (9 gray), bone marrow transplantation (BMT) was performed within 24 h after irradiation. One month after BMT, recombinant mouse SCF (200μg/kg/day, diluted in saline) (PeproTech, Rocky Hill, NJ, USA) and recombinant human G-CSF (50μg/kg/day, diluted in 5% dextrose) (Amgen, Thousand Oaks, CA, USA) or an equal volume of vehicle solution (50% of saline with 50% of dextrose) were subcutaneously injected for 5 consecutive days. Mice were sacrificed 6∼8 h after the final injection.

### 4.9. Bone Marrow Transplantation

UBC-GFP mice were deeply anesthetized. The femur and tibia bones were removed, cleaned of all connective tissue, and placed on ice in Hanks’ Balanced Salt Solution (HBSS). As described earlier, the ends of each tibia and femur bone were clipped to expose the marrow. The bone marrow was flushed out by ice-cold HBSS using a syringe with a 21-gauge needle. The bone marrow cells were filtered through a 70µm nylon mesh strainer (BD, USA), centrifuged at 300 g for 10 minutes at 4°C, and resuspended in ice-cold HBSS into a single cell suspension. Cells were transplanted to irradiated CCR5^-/-^ or WT mice within 24 h after irradiation (9 gray) by tail vein injection (1 x 10^7^ bone marrow cells in 0.7 ml HBSS per mouse).

### 4.10. Immunofluorescence staining

Mice were deeply anesthetized and perfused through the left cardiac ventricle with cold PBS followed by 10% phosphate buffered formalin (Fisher scientific, Waltham, MA, USA). Brains were removed and immersed in the same fixative overnight at 4°C and were then cryoprotected with 30% sucrose in PBS at 4°C for 2 days. Coronal brain sections, 30 µm thick, were cut with a cryostat (Leica Biosystems, Wetzlar, Germany). Free-floating technique was used for immunofluorescence staining. Briefly, brain sections (3-4 adjacent sections/brain) (1.10 mm posterior to bregma) were chosen for immunofluorescence staining. The sections were rinsed with PBS and incubated in a solution of PBS-diluted 10% normal goat serum or normal donkey serum (Jackson ImmunoResearch Laboratories, West Grove, PA, USA), containing 1% bovine serum albumin (BSA) (Sigma-Aldrich, St. Louis, MO, USA), and 0.3% Triton X-100 (Sigma-Aldrich, USA) for 1 hour at room temperature to block non-specific binding. The sections were then incubated with the primary antibodies including rabbit anti-cKit (1:100, Santa Cruz Biotechnology, Dallas, TX, USA), rabbit anti-GCSFR (1:100, Santa Cruz Biotechnology, USA), rat anti-CD31 (1:200, MEC 13.3, BD Pharmingen, San Diego, CA, USA), rabbit anti-CD31 (1:50, Abcam, Cambridge, UK), rabbit anti-Ionized calcium binding adaptor molecule 1 (Iba1, 1:400, Wako, Osaka, Japan), and goat anti-GFP (FITC-conjugated) (1:300, Abcam, UK) overnight at 4°C. The primary antibodies were diluted in PBS solution with 1% BSA and 0.3% Triton X-100 overnight at 4°C. In negative control brain sections, the primary antibodies were omitted. The following day, the sections were washed with PBS and incubated with the secondary antibodies diluted in PBS solution with 1% BSA and 0.3% Triton X-100 at room temperature for 2 h in the dark. The secondary antibodies used in this study were Cy2-conjugated goat anti-rat (1:200, Jackson ImmunoResearch laboratory, USA), Dylight-549-conjugated goat anti-rabbit (1:400, Jackson ImmunoResearch laboratory, USA), Alexa-Fluor 594-conjugated donkey anti-rabbit (1:500, Life Technologies, Carlsbad, CA, USA), and Alexa-Fluor 488-conjugated donkey anti-goat (1:500, Life Technologies, USA). Sections were rinsed with PBS again, mounted on Superfrost Plus Slides (ThermoFisher Scientific, Liverpool, NY, USA) and coverslipped with Vectashield Antifade Mounting Medium (Vector Laboratories, Newark, CA, USA). Z-stack images of 15 layers with an interval of 1.0μm in the cortex were taken with an LSM 780 confocal fluorescence microscope (Zeiss, Oberkochen, Germany). Bone marrow-derived cells adhering to the endothelial cells were determined by the co-localization of the GFP^+^ cells and CD31^+^ cells under the orthogonal view using ImageJ software. Monocytes/macrophages surrounding blood vessels were defined as GFP^+^/Iba1^+^ monocytes/macrophages within 10µm from CD31^+^ blood vessels. The number of GFP^+^ cells adhering to the CD31^+^endothelial cells, and the number of GFP^+^/Iba1^+^ monocytes/macrophages surrounding the blood vessels were quantified.

In the *in vitro* experiment, bEnd.3 cells were fixed in 4% cold PFA (Polysciences, Inc. PA, USA), and the immunofluorescence staining was performed following the same procedures as stated above.

### 4.11. Statistical Analysis

Data analysis was performed in a blind manner. Two-tailed t-tests were used to determine significant differences between two groups. One-way ANOVA followed by *post-hoc* Tukey’s test was used for comparison of three or more groups. For all tests, probability values < 0.05 were considered statistically significant. Data are presented as mean ± SEM.

## Supplementary Materials

The supporting information including Figures S1 and S2 can be downloaded.

## Author Contributions

Conceptualization, L.R.Z., X.F.R.; methodology, X.F.R., J.C.H., and H.H.; formal analysis, X.F.R., J.C.H.; investigation, X.F.R., J.C.H.; resources, L.R.Z., S.K.; data curation, X.F.R., J.C.H.; writing—first draft preparation, X.F.R., J.C.H.; writing—review and editing, L.R.Z.; visualization, X.F.R., J.C.H.; supervision, L.R.Z.; project administration and funding acquisition, L.R.Z. All authors read and approved the final version of the manuscript.

## Institutional Review Board Statement

All experiments were carried out in this study in accordance with protocols approved by Institutional Animal Care and Use Committees. The protocol (Approval Code: P-09-024; Approval Date: February 17, 2009) was approved by Louisiana State University Health Sciences Center Institutional Animal Care and Use Committee. The protocol (Approval Code: 369; Approval Date: August 2, 2016) was approved by the State University of New York Upstate Medical University Institutional Animal Care and Use Committee.

## Informed Consent Statement

Not applicable.

## Availability of data and materials

The datasets generated during the current study are available from the corresponding author on reasonable request.

## Acknowledgments

Authors would like to thank Mrs. Michele Kyle and Karen Hughes for their assistance in proofreading the final version of this manuscript.

