## Supplementary Figure Legends for "Hematopoietic growth factors Regulate Entry of Monocytes into the Adult Brain via Chemokine Receptor CCR5"

**Supplementary Figure 1.** SCF in combination with G-CSF enhances Iba-1^+^ monocyte adhesion to endothelial cells. (**A**) Quantification data. Mouse brain-derived endothelial cells (i.e. bEnd.3 cells) were incubated with medium alone (con: control), SCF+G-CSF (20ng/ml), TNF-α (100ng/ml), and LPS (1µg/ml) for 16-18 hours. After washing, Iba-1-GFP^+^ monocytes were added to the bEnd.3 cells. The monocyte-endothelial cell adhesion assay was performed using flow cytometry. Mean ± SEM. Repeated by 3 independent experiments. ***p* < 0.01, ****p* < 0.001 *vs.* the medium control. One-way ANOVA followed by *post-hoc* Tukey’s test. (**B**) Representative dot plots of flow cytometry showing adhesion of Iba-1-GFP^+^ monocytes to bEnd.3 cells that were pre-treated with medium alone, SCF+G-CSF (20ng/ml), TNF-α (100ng/ml), and LPS (1µg/ml) for 16-18 hours.

**Supplementary Figure 2.** The mRNA expressions of other chemokine receptors on bEnd.3 cells are not changed by SCF+G-CSF treatment. (**A-F**) Quantitative real-time PCR data. The mRNA expressions of CCR1 (**A**), CCR2 (**B**), CCR3 (**C**), CCR4 (**D**), CCR8 (**E**), and CXCR4 (**F**) in bEnd.3 cells that were cultured for different time periods (30min to 6 hours) in the presence of medium alone (con: control), SCF+G-CSF (20ng/ml), TNF-α (100ng/ml), and LPS (1µg/ml). Mean ± SEM. Repeated by 3 independent experiments. ***p* < 0.01, ****P* < 0.001 *vs.* medium control. One-way ANOVA followed by *post-hoc* Tukey’s test. nd: not detected.
