## Supplementary figures and images for "Hematopoietic growth factors Regulate Entry of Monocytes into the Adult Brain via Chemokine Receptor CCR5"

### Supplementary Figure 1

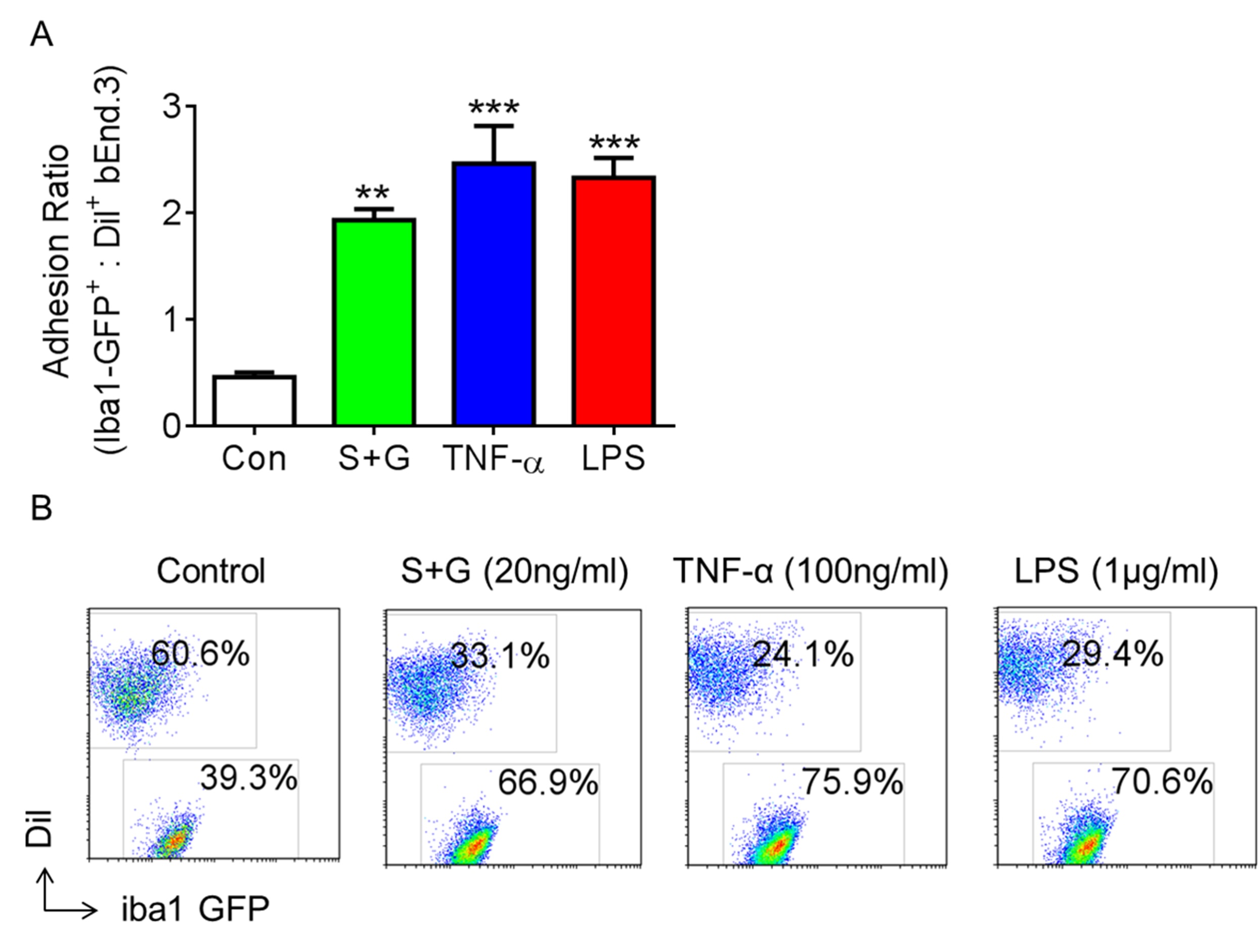

### Supplementary Figure 2

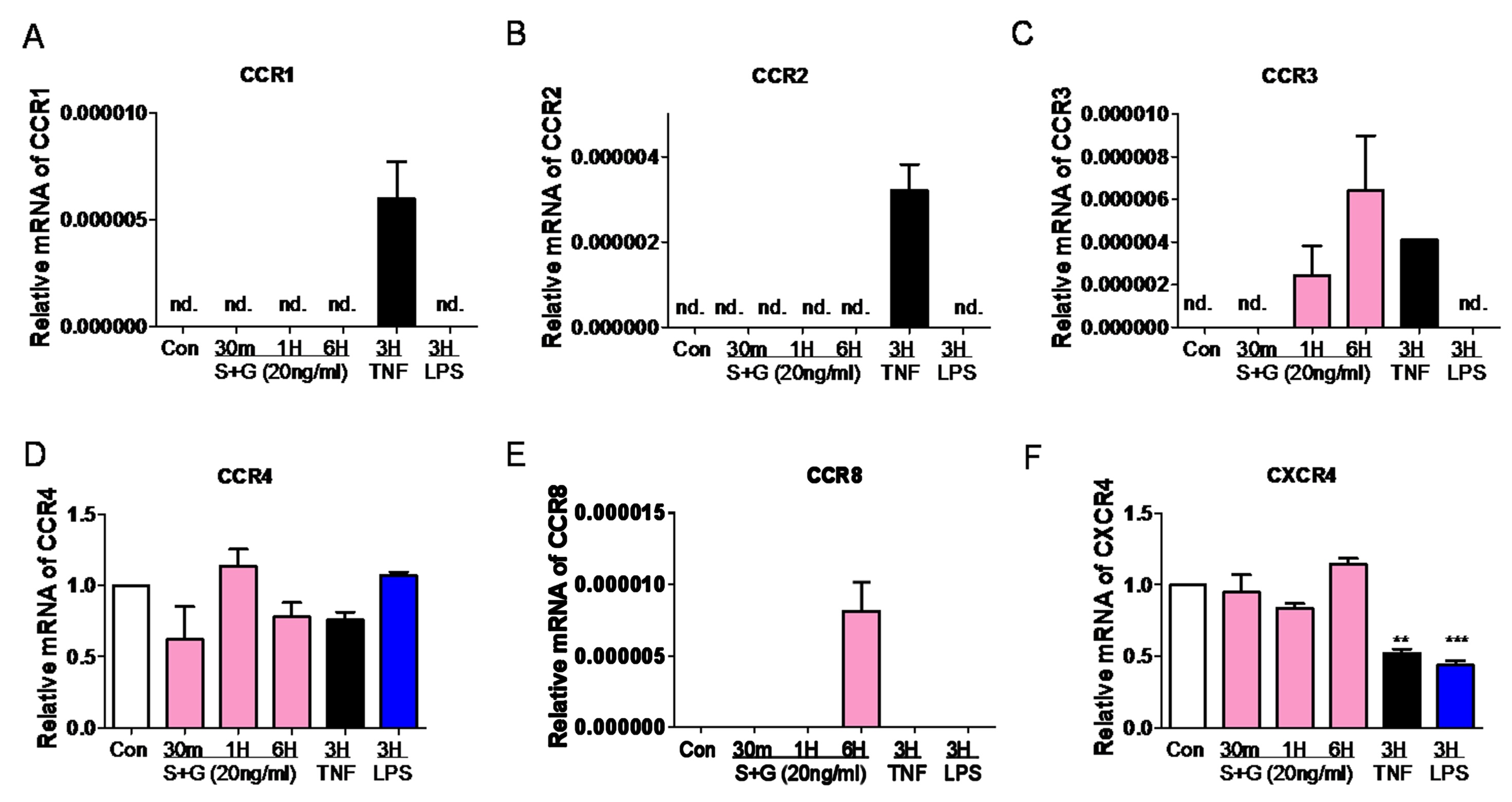
